# Fish microbiota repels ovipositing mosquitoes

**DOI:** 10.1101/2023.02.15.528615

**Authors:** Nimrod Shteindel, Yoram Gerchman, Alon Silberbush

## Abstract

The mere presence of predators causes prey organisms to enact predation-avoidance strategies. This presence is often reveled through predator-released kairomones. It was previously suggested that in many cases, the predator’s microbiota composition plays an important role in the release of these kairomones, however this mechanism is still poorly understood. Ovipositing mosquito females of several species are repelled by kairomones released from larvivorous fish. In this study we looked into the effects of the microbiota originated by *Gambusia affinis* (Baird and Girard) on the ovipisition behavior of gravid mosquito females in an outdoor mesocosm experiment. We show that interference with the fish microbiota significantly reduces its repellant effect. We further show that the bacterium *Pantoea pleuroti* isolated from the skin of the fish repels oviposition of *Culex laticinctus* (Edwards) and *Culiseta longiareolata* Macquart mosquitoes similarly to the effect of live fish. These results highlight the importance of bacteria in the interspecies interactions of their hosts and the potential conflict of interests in this system, where bacteria may benefit from the absence of the bacterivore mosquito larvae, but the fish loose access to prey.

## Introduction

During habitat selection, an individual evaluates numerous factors that will ultimately affect its decision. These, often conflicting, factors include the location, availability and quality of food resources, presence of a mating partner, competition and predation, which is one of the most important forces shaping prey organism decisions (Abramsky et al. 1990, Morris 1988, Rosenzweig 1981). The outcome of failure in predator detection and predation avoidance carry the most unforgiving ramifications to an individual’s fitness (Chesson and Kuang 2008, Lima and Dill 1990, Vamosi 2005), making the mere nonlethal presence of a predator a critical factor in determining the quality of a habitat (Brown and Kotler 2004, Hernández and Laundré 2005, Peacor and Werner 2001). In freshwater systems fish are considered the most effective predators with the highest impact on aquatic communities (Wellborn et al. 1996).

Prey organisms can detect fish using specific chemical signals released by the fish (Pohnert et al. 2007, Weiss et al. 2012). Fish-released kairomones are well known to induce morphological defenses in freshwater crustaceans (Tollrian and Dodson 1999, Weiss et al. 2012), which also use these signals to avoid habitats with high densities of predatory fish (Araújo et al. 2020, Decaestecker et al. 2002, Weiss et al. 2012). Aquatic amphibians and insects (e.g. beetles and dragonflies also use fish kairomones in selecting oviposition site. Such detection can produce a dramatic benefit – dramatically lowering larvae predation (Binckley and Resetarits Jr. 2003, Resetarits Jr. and Wilbur 1989).

This strategy was studies in mosquitoes of the *Culex* genus, reacting to the presence of the mosquito fish - *Gambusia affinis* (Cyprinodontiformes: Poeciliidae), known as a particularly effective predator of mosquito immatures (Becker et al. 2010, Blaustein 1992, Vinogradova 2000). Ovipositing *Culex* female detect and avoid pools containing *G. affinis* using fish-released kairomones (Angelon and Petranka 2002, Eveland et al. 2016). This repellent affect is specific to this fish species by comparison to other larvivorous fish (Silberbush and Resetarits Jr. 2017, Cohen and Silberbush 2021).

Although fish-released kairomones are known to be important in shaping of aquatic communities and ecosystems (Werner and Peacor 2003, Wellborn et al. 1996, Weiss et al. 2012) surprisingly little is known about them. Several studies suggested that fish-released kairomones are associated with fish microbiota (Akkas et al. 2010, Beklioglu, Telli et al. 2006, Beklioglu, Cetin et al. 2006, Landeira-Dabarca et al. 2019, Ringelberg and Van Gool 1998) yet no specific bacteria were identified. Here we studied the effect of Gambusia fish *Gambusia affinis* (Baird and Girard) skin microbiota on oviposition behavior of gravid females from the *Culex* genus. We tested the effect of antimicrobial substances applied to the fish and the effect of specific bacteria isolated from the fish skin on mosquito oviposition site choice in mesocosm experiments. Results show that interference with fish microbiota reduces their repellant effect and that specific bacterial isolates produce a repellant effect similar to that of fish.

## Methods

### Fish

*Gambusia affinis* fish were bought from local aquarium store and kept in an outdoor 1200 L holding tank (~100 fish). Fish were returned to the tank after each experiment.

### CuSO_4_ treatment of fish

Ten fish were placed in a 20 L container with a low (1 mg/L in aged tap water) CuSO_4_ solution at room temperature for 24 hours. This concentration was assumed to be well beneath the lethal concentration for this fish (Wallen et al. 1957). After the treatment fish were washed from the residual copper by net-transfer to 20 L of aged tap water, and left there for 24 hours before use for oviposition experiments. Untreated fish were simultaneously kept in tap water and treated as above.

### Isolation and identification of bacteria from fish skin

Treated and untreated *Gambusia affinis* (10 fish per treatment) were sampled for bacteria by swabbing their skin with sterile Q-tip prewetted with solution containing 0.9% NaCl and 0.02% Tween 80. The swabs were smeared on Tryptic Soy Agar plates (TSA; Himedia, India) that the plates incubated at 28 °C overnight. Morphologically represented colonies were isolated 5 times on TSA until uniform morphology was evident. The 16S RNA gene of selected colonies that were present in untreated fish but absent from treated fish (following visual morphological assessment) were amplified by PCR using the primer pair 11F (5’-GGATCCAGACTT TGATYMTGGCTCAG-3’) and 1512R (5’-GTGAAGCTTACGG(C/T)TAGCTTGTTACGACTT-3’) (Felske et al. 1997). Amplicons were sequenced by McLabs (USA) using the same primers and sequencing results identified using the EZBiocloud database (Yoon et al. 2017). Complete list of isolated and identified bacteria species is found in Table 1.

**Table 1:**
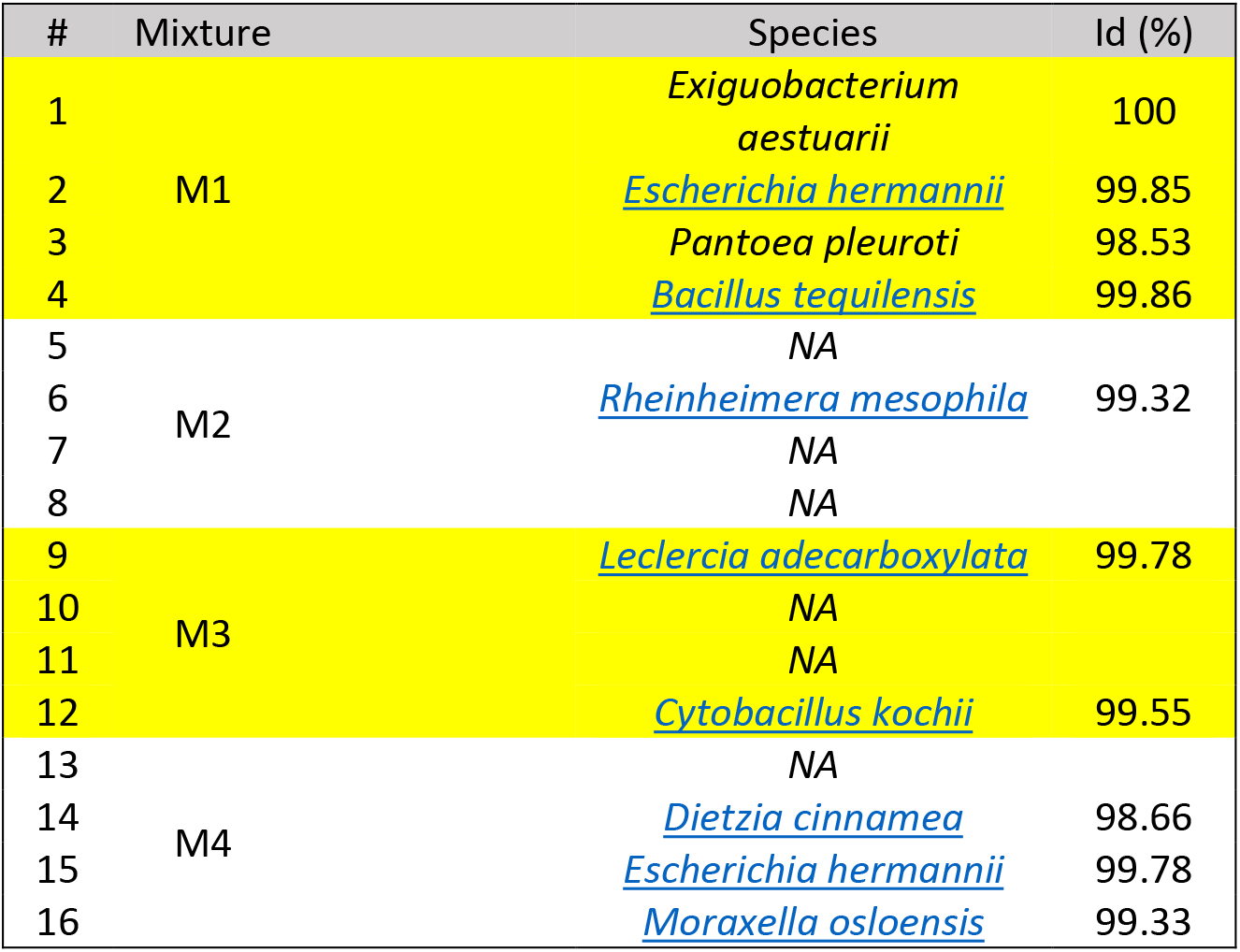
Bacterial isolates from untreated fish skin.

### Quantification of oviposition

Oviposition was quantified in a field experiment using pool mesocosms. Each pool was a black plastic pan (66.04 × 50.8 × 15.24 cm) containing 50-L of aged tap water. Pools were located in the botanic garden of Oranim college, Kiryat Tivon, Israel, organized in blocks, each block containing one pool per experimental condition. Egg rafts were counted and removed daily until the experiment was terminated. Egg rafts were placed in plastic jars and allowed to hatch and larvae identified to species after developing to fourth instar using (Becker et al. 2010).

First experiment – The effect of altered fish microbiome on oviposition: We placed live fish inside plastic cylindrical cages (22 cm diameter) with multiple one mm diameter holes and openings covered with a fiberglass screen. Cages were either empty fishless controls, or included 1 single fish. Ten grams of rodent pellets (Ribos rodent pellets-17% protein) were added to each pool outside the fish cages to encourage oviposition. In all cases treatments were randomly assigned within each block of adjacent pools. Fish in the experimental field pools were not fed and replaced every 3 days. Treatments within each block were the following: 1) Fishless control (empty cages), 2) untreated fish, and 3) treated fish-pre-treated with CuSO_4_ to remove bacteria and washed thoroughly.

Second experiment – Effect of specific bacteria mix on oviposition: In the second field experiment we tested the effects of specific bacteria isolated from *G. affinis* and demonstrating difference between CuSO_4_ treated and untreated fish. To simplify the experiments, it was done in mixes of four isolates each (1-4, 5-8, 9-12, and 13-16 out of total 16 isolates, see table 1). Each bacteria isolate was cultivated separately in tryptic soy broth (TSB) medium, diluted to OD_600nm_ = 1 in TSB and further diluted to a 50 L pool. Each isolate was grown separately in TSB overnight, and 12.5 mL of each isolate added to a 50 L pool. Control pools were treated with 50 ml fresh TSB.

Third experiment – Effect of specific bacterial isolates on oviposition: In the 3^rd^ experiment, isolates 1 to 4, found the most influential in the 2^nd^ experiment, were grown as above, diluted in TSB to OD_600nm_ = 1, and 50 ml of each isolate diluted into each 50 L pool, resulting in Final OD_600nm_ ~0.001, corresponding to ~10^5^ cells/mL, well within the range of bacterial cell densities to be expected in a ‘dirty’ pond (Ibekwe et al. 2007). Control pools were treated with 50 ml fresh TSB. Pools were set in 6 blocks of 5 treatments each – one of each four isolates and TSB as control.

### Statistical analysis

We used the total, per pool, number of all egg rafts collected for each mosquito species across all dates as a dependent variable. We used square root transformations of these values with an addition of 0.5 to all values to homogenize among-treatment variance (Yamamura 1999). Homogeneity of variance was tested using Levene’s test. We conducted separate univariate ANOVAs for each mosquito species in each of the experiments using “Block” and “Treatment” as fixed factors. In cases when Block affect was not close to significance (p>0.25), it was rolled into the error term. Treatment means were compared using Fisher’s protected LSD, when the main effect of treatment had p<0.1 using α = 0.05 for individual LSD comparisons. All analyses used SPSS statistics for windows version 24 (IBM 2016).

## Results

Of the total egg rafts sample, 42% were identified as *Culex laticinctus* Edwards, and 30% as *Culiseta longiareolata* Macquart. The remaining 28% egg rafts were identified as *Culex pipiens* Linnaeus and *Culex perexiguus* Theobald. The latter two species was excluded from analyses due to low densities.

First field experiment: The number of egg rafts of both *C. longiareolata* and *C. laticinctus* varied significantly among the treatments with number of egg rafts in pools with untreated fish significantly lower in comparison to fish-less control pools (Figure 1). This trend was not observed for pools containing treated fish (F_2,12_=4; p=0.05 and F_2,12_=6.4; p=0.013 for the two species respectively).

**Figure 1.**
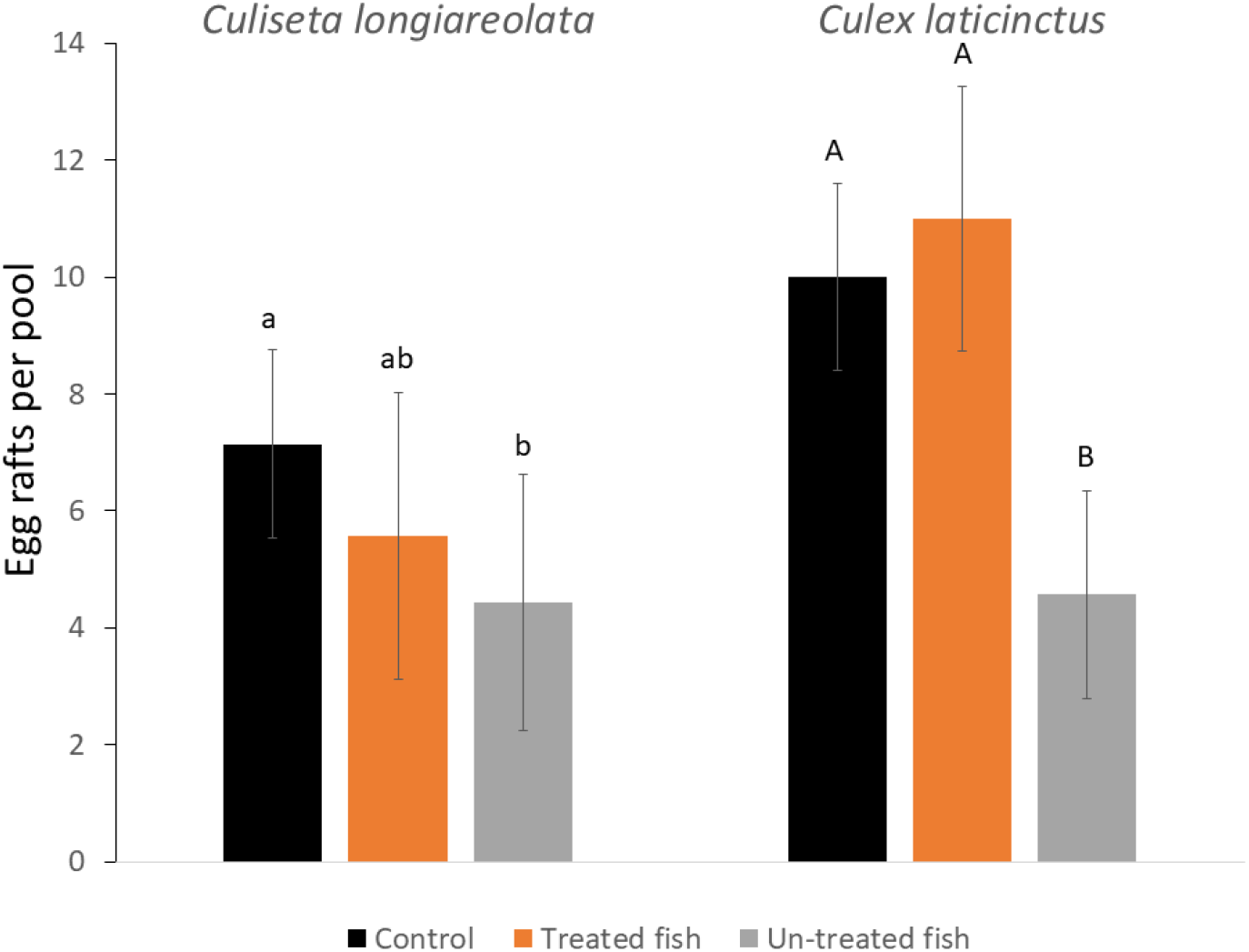
Oviposition of *Culiseta longiareolata* and *Culex laticinctus* Numbers are mean number of egg rafts per pool (±1 SE). Letters indicate treatments that are significantly different.

Second field experiment: Egg raft distribution of *C. longiareolata* did not vary among the different treatments (F_4,20_=1.7; p=0.2). By contrast, the distribution of *C. laticinctus* egg rafts was affected by the different treatments (F_4,20_=2.8; p=0.05). Pools containing the mixture of the first 4 isolates (M1; table 1) received significantly less egg rafts in comparison to control pools (Figure 2).

**Figure 2.**
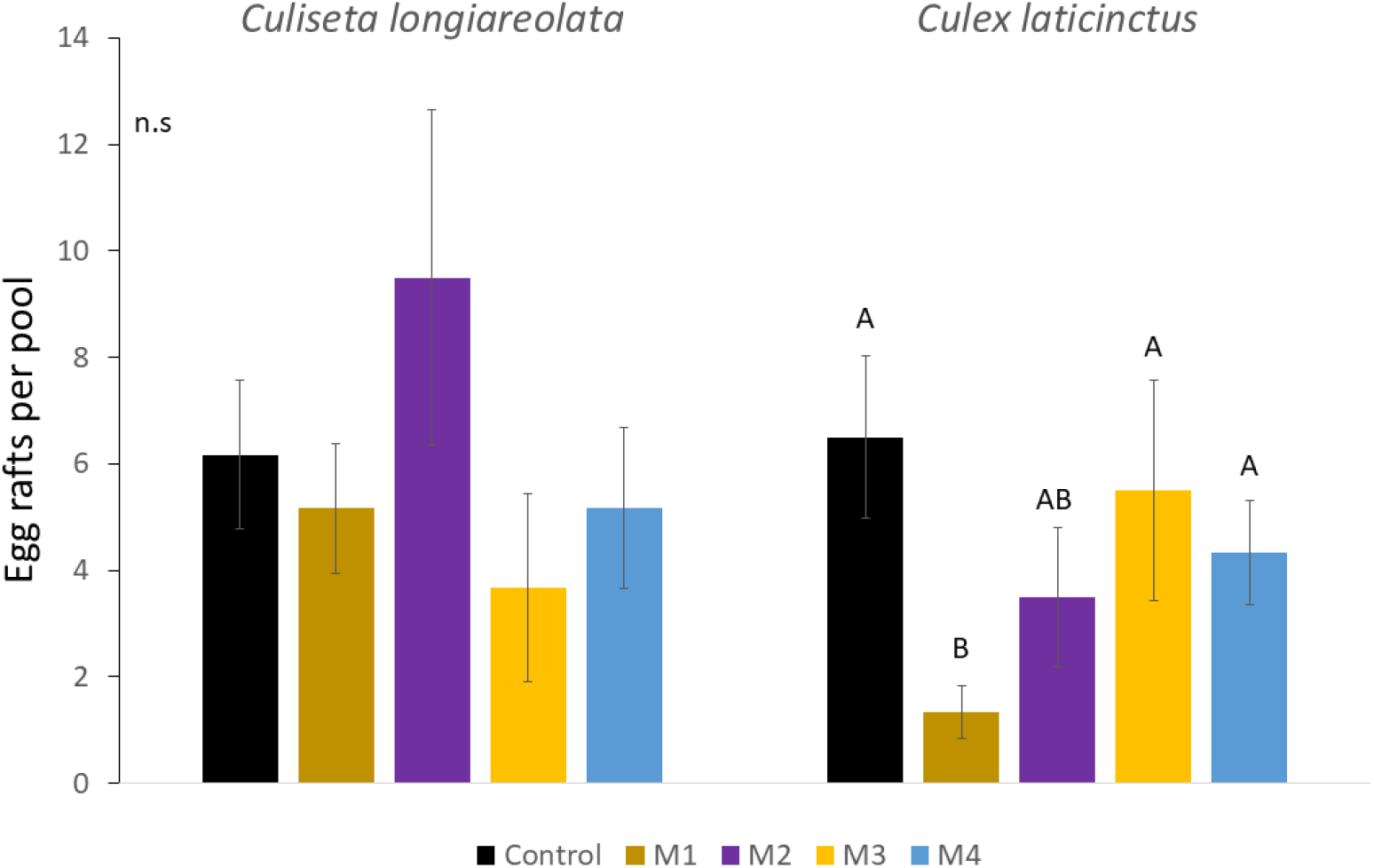
Oviposition of *Culiseta longiareolata* and *Culex laticinctus* Numbers are mean number of egg rafts per pool (±1 SE). Letters indicate treatments that are significantly different.

Third field experiment: *Culiseta longiareolata* egg rafts varied significantly among the treatments (F_4,20_=8.5; p<0.001). Pools treated with *Exiguobacterium aestuarii* received significantly more egg rafts in comparison to control. By contrast, pools treated with *Bacillus tequilensis* or *Pantoea pleuroti* received significantly less egg rafts in comparison to control. *Culex laticinctus* egg rafts distribution likewise varied among the treatments (F_4,25_=2.6; p=0.06) with pools treated with *P. pleuroti* receiving significantly less egg rafts in comparison to control pools (Figure 3).

**Figure 3.**
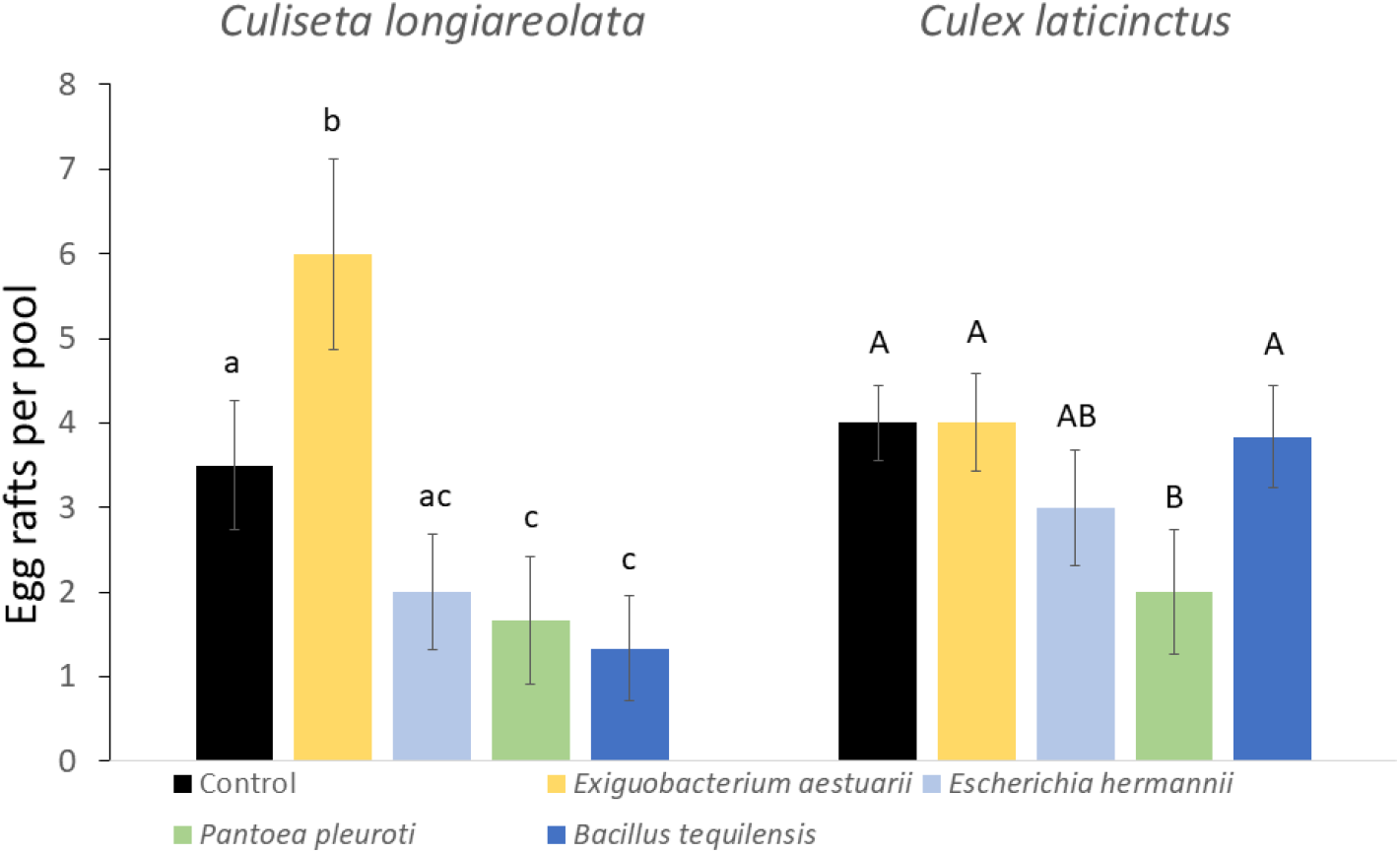
Oviposition of *Culiseta longiareolata* and *Culex laticinctus* Numbers are mean number of egg rafts per pool (±1 SE). Letters indicate treatments that are significantly different.

## Discussion

Chemical signals produced by associated microorganisms have been shown to play an important role in animal communication (Ezenwa and Williams 2014). This phenomenon is known mostly in the context of pheromones production, but also take part in intra-species interaction. Microbiota-generated volatiles are exploited by eavesdropping parasites or predators who use them as kairomones to detect the presence of host or prey (Mazorra-Alonso et al. 2021). Some studies suggested that these volatiles can also be used by prey species to detect predator, for example, larval mayflies show anti-predator behavior when exposed to hexomines produced by fish mucus dwelling bacteria (Landeira-Dabarca et al. 2019). Ovipositing *Culex* female are repelled by the presence of *G. affinis*, even when the fish are caged or previously removed (Angelon and Petranka 2002, Eveland et al. 2016), indicating that the source of this effect is fish-released kairomones. It was previously suggested that these kairomones may originate from bacterial symbionts associated with *G. affinis* (Why and Walton 2020, Why et al. 2021), however this is the first study showing a specific bacterium used by prey to detect and avoid predator presence. *Gambusia affinis* repellent effect was reduced dramatically when fish were treated with an antibacterial substance, suggesting that skin bacteria are the source of these kairomones (Figure 1). This effect did not fade over time, indicating that the bacteria originate from the fish casings. Specific odors may indicate specific bacterial communities associated with different fish (Mazorra-Alonso et al. 2021), explaining the different responses of the mosquito to different fish species (Silberbush and Resetarits Jr. 2017, Cohen and Silberbush 2021) and fish chemical camouflage towards ovipositing treefrogs and colonizing beetles (Resetarits Jr. and Binckley 2013).

The results presented here raise interesting questions regarding the definition of a kairomone, defined as a trans-specific chemical messenger the adaptive benefit of which falls on the recipient rather than on the emitter (Brown Jr et al. 1970). If we consider the fish and its bacterial symbioses as a holobiont, the signals originating from the fish skin microbiota are indeed kairomones, but this perspective assumes a certain unity of interests between the fish and the bacteria. In this case however, the fish microbiota repels gravid mosquitos and reduces the number of mosquito larvae, which serves as prey for the fish, but predators of bacteria. In this case of “life vs. dinner”, the “life” reward to both mosquito and bacteria outweighs the benefit of the fish dinner in the dynamics of the system. This merits a future study to quantifies the costs and benefits in the system – the potential benefits to the fish from its natural microbiota, the predation risk to the specific symbioses from the mosquito larvae and the impact of reduced prey availability of the fish fitness. Furthermore, the data suggest that wise use of bacteria, tailored to repel or attract specific mosquito species while minimizing the effect on harmless and potentially important species, could result in decrease in mosquito-born disease spread.

